# Expanding the SEVA plasmid toolbox for tuneable control of gene expression in *Synechocystis* sp. PCC 6803

**DOI:** 10.64898/2026.11.16.767122

**Authors:** Govinda Guevara, Juana María Navarro Llorens, Arantxa Marco, Sara Baldanta

## Abstract

The use of cyanobacteria as biotechnological chasses is considered a key point in the circular economy due to their scarce growth requirements and its capacity of fixing atmospheric CO_2_. However, for obtaining the most of their potential, it is often necessary to count with tools that allow the modification of the genetic expression.

In this work, we have characterized a molecular toolbox for reliable control of gene expression control through induction or repression in *Synechocystis* sp. PCC6803 using SEVA plasmids. Different inducible promoters showing a variety of dynamic range, strength and leakiness using GFP our YFP as reporter genes were assayed.

On the other hand, we also evaluated the gene repression in cyanobacteria by using CRISPRi technology, which allows to repress the gene transcription in a targeted manner. Employing an inducible promoter, a new SEVA-dCas9 based vector was built for *Synechocystis* sp. PCC6803. The vector contains dCas9 (DNase-dead Cas9 nuclease) under the tightly regulated PBADRSW promoter, which combines a theophylline-responsive riboswitch with a rhamnose-inducible promoter. The sgRNA expression cassette is also encoded on the same plasmid, resulting in a compact, self-contained CRISPRi system that simplifies genetic implementation and enables targeted gene. As a proof of concept, we demonstrated that the system enables reversible knockdown of a chromosomally integrated eYFP reporter driven by the constitutive Pc223 promoter in *Synechocystis* sp. PCC 6803.

These results provide new insights into the genomic strategies and available promoters and CRISPRi-SEVA plasmids for the industrial cyanobacteria *Synechocystis* sp. PCC 6803 that would be valuable for further applications.

## Introduction

Cyanobacteria are photosynthetic microorganisms capable of fixing atmospheric carbon dioxide, playing a fundamental role in global carbon cycling and Earth’s ecosystems. Beyond their ecological importance, they have emerged as promising biotechnological chassis for the sustainable production of a wide range of valuable compounds (Lindberg et al., 2023; Liu et al., 2024; Angeles et al., 2025; Bolay et al., 2025; Chen et al., 2025; Gallego et al., 2025; Walshe and Saha, 2026). Their potential as microbial cell factories stems from several key advantages: i) they do not require organic carbon sources for photoautotrophic growth; ii) they contribute to atmospheric carbon dioxide fixation; iii) their cultivation requirements are quite basic: mainly light energy, water and carbon dioxide and, iv) many species are amenable to genetic manipulation. However, there are still key limitations to engineered pathway production, including metabolic burden, polyploidy, slow photoautotrophic growth in some strains, native regulatory constraints, and cultivation challenges that restrict productivity and yields (Walshe and Saha, 2026).

In fact, production of various chemicals has been reported but most of them are proof-of-concept-level at a low product yield and their largescale application is still technically and economically limited (Liu et al., 2024). Therefore, a big breakthrough in terms of efficiency and productivity is still demanded before they become industrial workhorses and entry into the commercial-scale production process (Lindberg et al., 2023; Chen et al., 2025; Zhang et al., 2025).

In the last years, there have been several attempts to improve the cellular robustness of cyanobacteria. At one hand, by facing environmental stresses through culture conditions: i) adaptive evolutionary strategy and, ii) semicontinuous algal culture supported by machine learning and synthesis (Liu et al., 2024; Murakami et al., 2026). On the other hand, the strategies for improving the efficiency of engineered cyanobacteria from both genetic and metabolic dimensions using synthetic biology include: i) altering endogenous gene expression by introducing heterologous genes or metabolic pathways ii) the use of artificial-intelligence-aided design methods for genetic elements such as the use of deep generative models using big data integration and, iii) the use of artificial intelligence and mathematical models to identify and optimize metabolic stuck points (Seo et al., 2023; Zhang et al., 2023; Liu et al., 2024; Kamalesan et al., 2026; Walshe and Saha, 2026). Several synthetic biology approaches have been developed and applied to cyanobacteria, including modular cloning systems such as CyanoGate Modular and CyanoConstruct Cloning (Vasudevan et al., 2019; Venero et al., 2026), the Start-Stop Assembly method (Taylor et al., 2021), recombineering techniques (Jones et al., 2021; Li et al., 2026), the development of synthetic genetic circuits (Taton et al., 2017; Zhang et al., 2022) and CRISPR-based genome engineering approaches (Yao et al., 2020; Baldanta et al., 2022; Cengic et al., 2022; Wang et al., 2023; Walshe and Saha, 2026).

Most current metabolic engineering and synthetic biology efforts in cyanobacteria have focused on a limited number of model strains, particularly from genus *Nostoc*, *Synechococcus* and *Synechocystis*, owing to their comparatively well-characterized biological and genetic backgrounds. These cyanobacteria possess several desirable features for their development as biotechnological chassis, including relatively small genomes, metabolic flexibility that allows growth under different nutritional regimes (photoautotrophic, mixotrophic and, in some cases, heterotrophic conditions), well-characterized genetic backgrounds, relatively fast growth rates, and amenability to genetic manipulation (Mills et al., 2020; Lindberg et al., 2023; Liu et al., 2024). Nevertheless, the genetic engineering toolbox available for cyanobacteria remains comparatively limited when compared with those established for model microorganisms such as *Escherichia coli*, *Pseudomonas putida* and yeasts. Moreover, it takes longer times to construct an engineered strain of cyanobacteria, their biological parts are not well-characterized, and there is still a lack of successful procedures for some non-model cyanobacteria (Bourgade and Stensjö, 2022; Cheng et al., 2023; Liu et al., 2024). Addressing these limitations requires the development and characterization of standardized genetic parts to expand the synthetic biology toolbox available for cyanobacteria and unlock their potential as next-generation microbial cell factories.

Promoters are key genetic elements for controlling gene expression at the transcriptional level, making them essential components for the design and optimization of synthetic genetic circuits and engineered metabolic pathways. Several promoters have been characterized in the model strain *Synechocystis* sp. PCC 6803 (hereinafter Syn6803), including both for constitutive (Ferreira et al., 2018; Wang et al., 2018; Vasudevan et al., 2019; Behle et al., 2020) and inducible expression (Englund et al., 2016; Behle et al., 2020; Seo et al., 2023). The latter include promoters regulated by different chemical and environmental stimuli, such as IPTG (Albers et al., 2015; Markley et al., 2015; Ferreira et al., 2018), metal ion (Englund et al., 2016; Pérez et al., 2017), light (Albers and Peebles, 2017; Das and Maiti, 2024), rhamnose or vanillic acid (Kelly et al., 2018; Meyer et al., 2019).

Other way to control gene expression is using the CRISPR Clustered Regularly Interspaced Short Palindromic Repeats) technologies, either using nucleases for gene editing or dead nucleases for transcription interference (CRISPRi). Both systems have been already implemented for *Syn6803* (Baldanta et al., 2022; Cengic et al., 2022; Miao et al., 2023; Veit et al., 2025; Enkerlin et al., 2026; Veit et al., 2026). CRISPR interference (CRISPRi) is a gene repression technology that relies on a catalytically inactive Cas protein (dCas), generated by thought point mutations that abolish its DNA-cleavage activity, and a single-guide RNA (sgRNA) designed to recognize a specific target DNA sequence. The dCas–sgRNA complex binds to the target DNA region adjacent to an appropriate protospacer-adjacent motif (PAM; 5′-NGG-3′ for *Streptococcus pyogenes* Cas9), thereby interfering with transcription and repressing target gene expression. dCas9-based CRISPRi enables targeted and multiplexed gene repression without requiring permanent changes to the genome. This approach has been used both to modulate the expression of multiple genes and to investigate the functions of genes that remain poorly characterized or unannotated (Kirtania et al., 2019; Miao et al., 2023; Hishida et al., 2024).

Despite the advances described above, there is still a need to expand and improve the genetic toolbox available for this industrially relevant cyanobacterium, particularly through the development of reliable tools that enable precise and rapid control of gene expression. To address this need, this study pursues two main objectives: first, to evaluate inducible promoters for *Synechocystis* sp. PCC 6803; and second, to extend the applicability of the previously established SEVA-based platform in cyanobacteria (Baldanta et al., 2022) by incorporating CRISPRi functionality, thereby providing a new tool for targeted gene repression in Syn6803. SEVA is a collection of modular vectors that enable the exchangeability of diverse elements (e.g. origins of replication and antibiotic selection markers) and the combination with many cargo sequences for varied end-applications (Martínez-García et al., 2023). Accordingly, several promoters were evaluated using a promoterless SEVA-based vector, and a new SEVA-based plasmid carrying the CRISPRi machinery was constructed for Syn6803.

## Material and Methods

### Bacterial strains and growth conditions

Cloning steps were carried out in *Escherichia coli* strain DH5α or DH10B while *Escherichia coli* strain HB101 was used for triparental conjugation. *E. coli* cells were grown at 37°C in liquid LB or on LB agar plates supplemented with 50μg/mL kanamycin, 10μg/mL chloramphenicol, or 50μg/mL spectinomycin as required.

Wild-type *Syn6803* was used as the control strain. Cells were grown in BG11 medium buffered to pH 7.5 with Hepes 10 mM on 1.5% agar plates or liquid medium at 30°C under 100μmol photon • m-2 • s-1 of continuous white light. Antibiotics spectinomycin 10μg/mL, chloramphenicol 10μg/mL and kanamycin 50μg/mL were utilized for the selection of cyanobacterial recombinant strains.

### Chemicals and reagents

All the inducers used were provided by Thermofisher. Rhamnose was prepared at 100g/L, IPTG at 1 M, ε-caprolactam at 1M and 3-methylbenzoate at 0.5M (all in water). Theophylline was prepared at 200mM in DMSO and incubated at 55°C until complete dissolution. T4 DNA ligase and restriction enzymes were purchased from Takara or ThermoFisher Scientific. KOD-Hot start polymerase (Novagen) or Phusion (Thermofisher) were used for high-fidelity polymerase chain reaction (PCR). For colony PCR, NZYTaq II 2x Green Master Mix was used. Minipreps and DNA purification kits were obtained from Nzytech or Thermofisher. Oligonucleotides (Table S1) and gBlocks (Table S2) were synthesized by Condalab and IDT. All protocols were conducted according to the manufacturer’s instructions. Plasmids maps were created with SnapGene 3.1 (GSL Biotech LLC).

### Construction of plasmids for promoter characterization

All the plasmids used in this work are listed in Table 1. As reporter genes, eYFP and msfGFP were used, hereafter named YFP and GFP respectively. To construct pSEVA251-P_c223_-YFP, P_c233_-YFP gBlock (Table S2) was amplified using SB72/69, digested with *Pml*I and *Pst*I and cloned in pSEVA251 previously digested with *Sma*I and *Pst*I. pALM174 and pALM179 plasmids were purchased from Addgene (Plasmid #82022 and Plasmid #82023 respectively). The cassette including the promoter, *yfp* and *lacI* was amplified by PCR with AL41/AL43 and then cloned into pSEVA251 using *Sal*I and *Spe*I giving pSEVA251-P_cLac94_-YFP and pSEVA251-P_cLac143_-YFP. pSEVA251-P_Pm-XylS_-GFP was constructed by inserting the *gfp* amplified by PCR with AL20/AL23 and digested with *Sac*I and *Pst*I in pSEVA258. For pSEVA251-P_nit_-GFP, a plasmid already constructed in the laboratory, was used to place the *gfp* under this P_nit_ promoter by amplifying the reporter with AL21/22 and cloning the product using *Nde*I and *Hind*III. Then, the cassette including promoter, *gfp* and *nitR* were transferred to pSEVA251 digesting the plasmid with *Kpn*I and *Spe*I, and *Kpn*I and *Xba*I for the insert. To construct pSEVA251-P_BAD_-GFP, a plasmid (pSB01) containing the sequence from gBlock P_BADRSW_ was used. GFP was amplified with AL38/39 from pSEVA247M, digested with *Sca*I and *Not*I and cloned in the plasmid pSB01 digested with *Eco*RV and *Not*I. Then, the whole cassette including also the double terminator B0015 and the *Km^R^* gene was amplified with SB67/68 and cloned into a pSEVA251 using *Pac*I and *Spe*I sites.

**Table 1.**
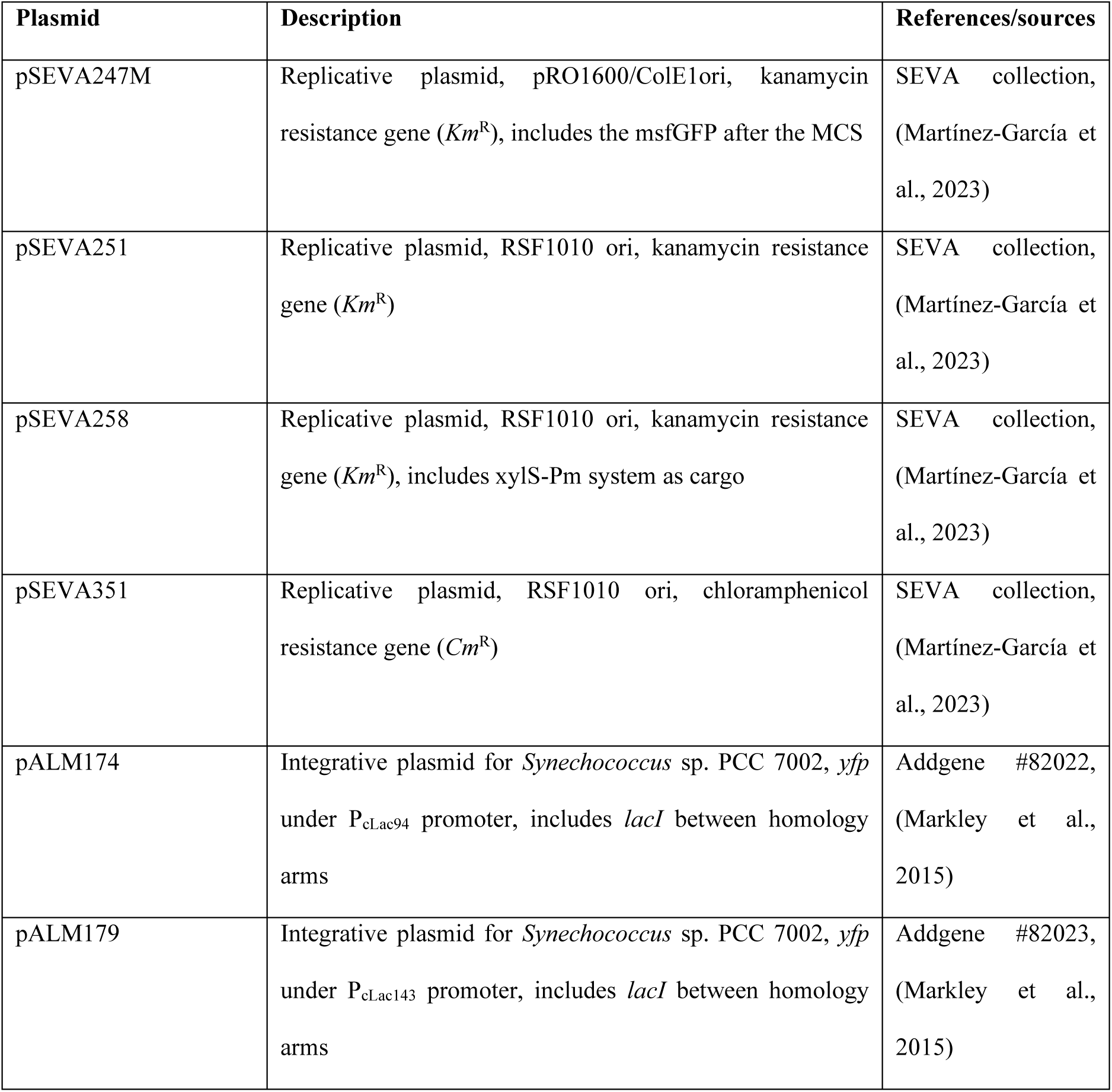
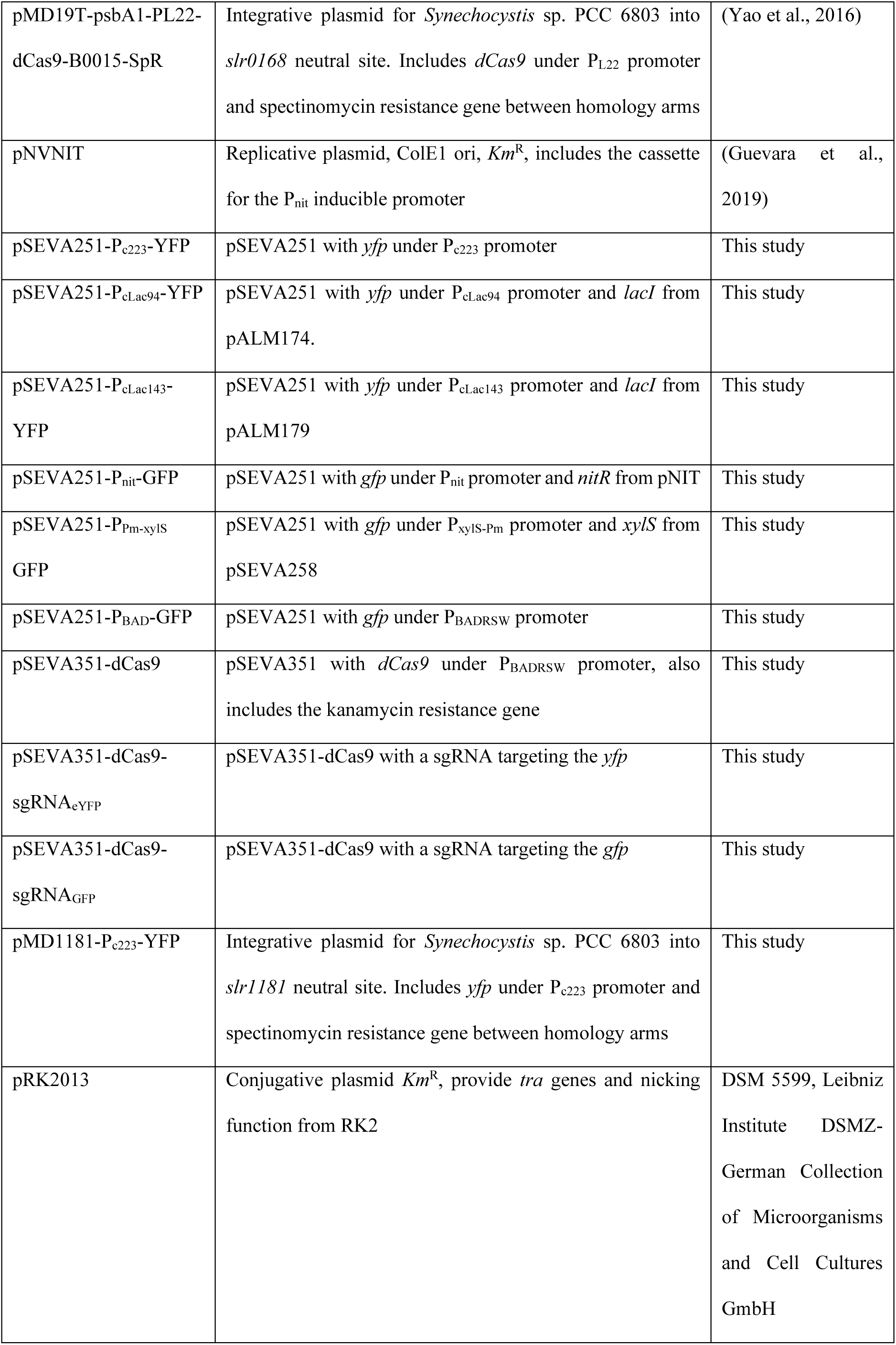
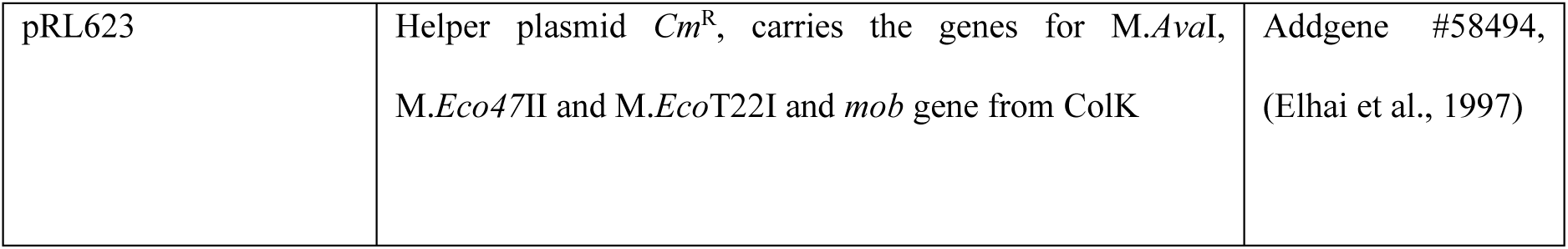
Plasmids used in this study.

### Construction of plasmids for CRISPRi-mediated repression

To test the CRISPRi in *Syn6803*, two plasmids were constructed. For YFP integration into the cyanobacteria genome, the YFP (with the terminator) under P_c223_ was digested from pSEVA251-P_c223_-YFP and cloned in pMD19T-psbA1-PL22-dCas9-B0015-SpR (Yao et al., 2016) using *Stu*I and *Eco*RI, yielding the integrative plasmid pMD1181-P_c223_-YFP. On the other hand, dCas9 was amplified from pMD19T-psbA1-PL22-dCas9-B0015-SpR with SARA60/61 by PCR. Then, the amplified DNA was digested with *Sma*I and *Not*I and cloned in pSB01. After that, the whole cassette including the dCas9 under P_BADRSW_ and the double terminator B0015 and the *KmR* gene were amplified with SB67/69 and cloned in pSEVA 351 previously digested with *Pac*I and *Spe*I.

A sgRNA targeting YFP (sgRNA_YFP_) was designed using Benchling (https://benchling.com.), with “single guide”, “20 guide length” and “PAM NGG SpCas9” as options. Genome GCA_0000097251 from *Syn6803* was used to calculate the possible off targets. The guide sequence selected was 32 nt downstream from +1 (TSS), targeted the non-template strand and had a high “off-target score” (meaning low off-targets expected (Hsu et al., 2013)). After the guide sequence, the sequence recognized by dCas9, and the *S. pyogenes* terminator were included, giving the sgRNA_YFP_. The sgRNA cassette (gBlock, Table S2) expressed the designed guide sequence under P_BADRSW_ promoter and was cloned between *Pac*I and *Eco*RI in pSEVA351-dCas9. To construct a control for CRISPRi repression, a sgRNA targeting GFP was designed (the same way as for *yfp*) and placed under P_BADRSW_ promoter using an overlap PCR with the primers AL50/48 and AL47/AL49, using the sgRNA_YFP_ as template. The overlap PCR replaces the sgRNA_YFP_ for a sequence targeting the GFP. The resulting sgRNA_GFP_ was cloned into pSEVA351-dCas9 using *Pac*I and *Eco*RI. Potential off targets in *yfp* were checked using Benchling.

The replicative plasmids, either pSEVA251 or 351-based, used for promoter characterization and CRISPRi were transformed into *Syn6803* by triparental mating, following the spot mating protocol (Elhai and Wolk, 1988). Briefly, *E.coli* HB101 strain bearing the conjugative plasmid pRK2013 was used as a conjugative strain. HB101 containing the replicative plasmid desired was the cargo strain. *Synechocystis* OD_750nm_ was adjusted to 0.1, mixed with *E. coli* and plated onto BG11 5% LB (vol/vol) agar. Plates were incubated for 48h at growth conditions. Conjugation filters were transferred onto BG11 agar supplemented with 50μg/mL kanamycin or 10μg/mL chloramphenicol. Colonies appeared within 7 days. To obtain the mutant strain with *yfp* under P_c223_ integrated at *slr1181*, natural transformation of *Syn6803* was performed as previously described (Baldanta et al., 2022) with the integrative vector pMD1181-P_c223_-YFP. Spectinomycin 10μg/mL was used for mutant colonies selection. Transformant colonies appeared after approximately 1 week. Up to 4 streaks were done before colony PCR was used for plasmid or genome integration validation. First fast DNA extraction with chloroform was done for each transformant and then 5 µL were used for PCR using NZYTaq II 2x Green Master Mix.

### Evaluation of promoters and CRISPRi in *Syn6803*

For the evaluation of constitutive or regulated promoters, *Synechocystis* colonies harboring the desired plasmid were grown in 3 mL of BG11 with the appropriate antibiotics. After 4 days, the cultures were diluted in fresh BG11 to a starting OD_750nm_ of 0.1 in 10 mL flasks. The cultures were grown for 24 hours and then, the inducers were added at final desired concentration: 1 mM IPTG, 20 mM ε-caprolactam, 1 mM 3-methylbenzoate, 2 mM theophylline and 2 g/L rhamnose.

For CRISPRi evaluation, cultures grew in 10 mL BG11 for a week and then diluted in fresh BG11 to a starting OD_750nm_ of 0.6 in 10 mL flasks. Two mM theophylline and 2 g/L rhamnose were added to induce the CRISPRi system. After 5 days, to evaluate the reversibility of YFP repression, 2 mL were washed 3 times and resuspended in fresh BG11 without inducers at an initial OD_750_ of 0.2. Samples were taken every 24 hours for fluorescence analysis until complete recovery of expression. Fluorescence was evaluated at different times for every condition, either for quantitative or visual analysis.

### Measurement of bulk fluorescence

Quantitative fluorescent analysis of the *Synechocystis* strains expressing either YFP and GFP for promoter characterization or CRISPRi assays was determined in a final 100 µL volume in 96-well black plates (Thermofisher) on a FLUOstar Omega plate reader (BMG Labtech). Cultures were adjusted to a OD_750nm_ of 0.1 and fluorescence measurements were normalized to this value. The excitation/emission wavelengths were set to 485/525 nm for GFP and 485/535 nm for YFP.

#### Microscopy visualization

Microscopy imaging was carried out on a Leica DFC345 FX fluorescence microscope using a 100× objective. At desired times of promoter or CRISPRi evaluation, 10 µL of culture were used for microscopy visualization. YFP was excited using a 485nm laser light and emission was collected at 525 nm. For chlorophyll visualization (autofluorescence), the excitation was performed at 577 nm and emission was collected at 603 nm. For each set of visualization, the gain was adjusted to avoid saturation of the higher signal employing the sample with the highest signal of fluorescence and then used the settled parameters for the rest of images, to ensure equal treatment.

## Results and Discussion

### Promoter evaluation in *Syn6803*

Synthetic biology applications often require well-characterized and strong promoters with predictable responses. The strength of promoters is generally determined according to the expression of a reporter gene such as GFP encoding green fluorescent protein or other fluorescence derivatives. Although there are several promoters reported for *Syn6803*, the reproducibility of the results is uncertain mainly due to external factors, for instance, the expression of a synthetic cassette being integrated into the genome may differ from being expressed within a vector. To improve reproducibility, a similar genetic backbone should be used. Even though, despite the use of the same regulatory system orthogonal PrhaBAD, and a high level of protocol standardization, up to ∼32% variation was found in promoter activity under induced conditions across different laboratories (Mager et al., 2023).

In order to evaluate different promoters in *Syn6803*, different steps were taken. First, a promoter-less vector was built using the replicative plasmid pSEVA251. This vector belongs to the SEVA repository, a collection particularly interesting owing to the modular and interchangeable structure of their plasmids, some of them such as pSEVA251 proved to replicate in cyanobacteria (Baldanta et al., 2022; Martínez-García et al., 2023). This vector contains the broad host-range RSF1010-ori that it has also been used in several plasmids to characterize orthogonal constitutive promoters in diverse cyanobacterial strains such as *Synechococcus elongatus* PCC 7942, *Syn6803*, *Anabaena* PCC 7120, *Leptolyngbya* BL0902 and the non-model extremophile *Nodosilinea* ML3B (Trieu et al., 2026).

On the other hand, promoters P_c223_, P_cLac94_ and P_cLac143_ previously tested in *Synechoccoccus* strains (Markley et al., 2015; Włodarczyk et al., 2020) were chosen for this study. P_c233_ is a constitutive promoter derived from *cpt* promoter (<u>c</u>yanobacterial <u>p</u>hycocyanin truncated promoter), which comes from *Syn6803cpcB* promoter. P_c233_ contains two mutations that increase its expression by two times as P_cpt_ and no significant difference is observed between high and low light conditions (Markley et al., 2015). On the other hand, cLac94 and cLac143 promoters are derived from P_cpt_ modified to be inducible by IPTG. The three promoters were cloned into the self-replicating vector pSEVA251 using the enhanced yellow fluorescent protein (eYFP) as reporter (Fig. 1a). Promoter constructs were transferred into *Synechocystis* by conjugation and an empty vector control was included. *Syn6803* cells carrying the respective promoter constructs were used to inoculate fresh cultures, which were grown for 24 hours before induction of expression by addition of IPTG if necessary. The cells were cultivated for an additional 72 hours, and the fluorescence from expressed eYFP was determined every 24 hours (Fig. 1b,1c). The cLac94 IPTG-induced promoter showed a low leakiness and higher induction rate, yielding around ∼80-fold increase of expression levels after 72 hours of IPTG addition. On the other hand, cLac143 was induced ∼25 times and had leaky expression without IPTG. Comparable results of fluorescence to cLac94 after 72 h postinduction were observed for the constitutive promoter P_c223_. Of the three validated promoters, cLac143 is the one that produced the greatest expression. The results were validated also by fluorescence microscopy (Fig 1d)

**Fig. 1.**
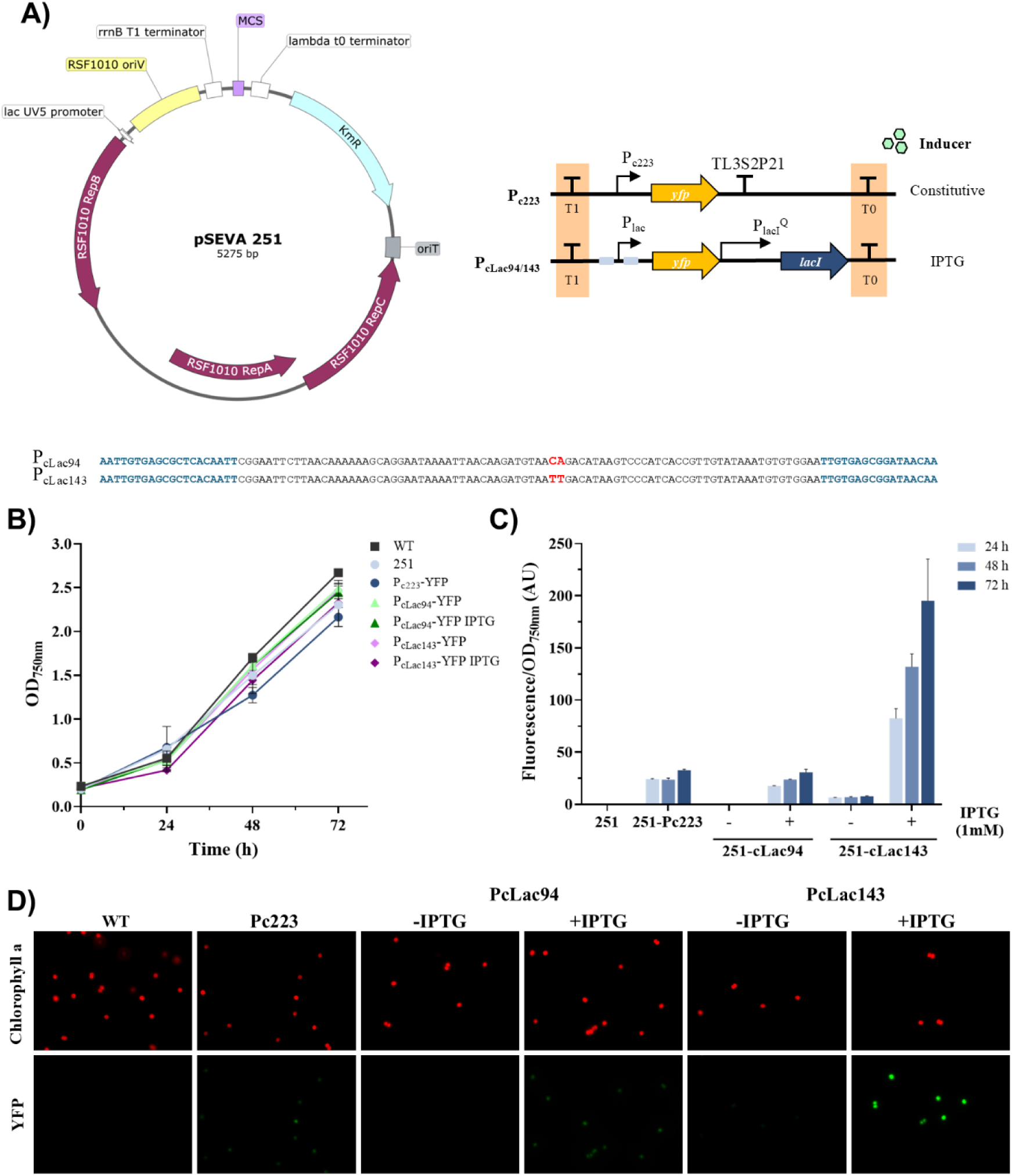
Promoter activities in *Syn6803*. **A)** A scheme of the pSEVA 251 plasmid core to evaluate promoters in *Syn6803*. The difference between both IPTG-inducible promoters is depicted in red. The *lac* operators are shown in blue. **B)** Growth curves of *Syn6803* recombinants cultured in BG11 medium. Error bars represent the standard deviations observed from at least two independent experiments. **C)** Fluorescence measurements from the different promoters. Activities were measured as eYFP fluorescence corrected by OD_750nm_ for cell cultures in BG11 medium. The results correspond to 24-, 48-and 72-hours post-induction. As control, Pc223 constitutive promoter was added. Error bars indicate standard deviations (*n*= 3). **D)** Fluorescent microscopy image of cells expressing eYFP driven by different promoters. Phase contrast, chlorophyll autofluorescence and eYFP fluorescence are shown.

Besides these three promoters, two inducible promoters from other microorganisms were chosen additionally (Fig S1). P_nit_ is a nitrile-inducible promoter from the actinobacteria *Rhodococcus rodochrous* (Pandey et al., 2009) and it has been used to control gene expression in mycobacteria and other Gram-positives. The expression cassette consists of the *nitR* gene (regulatory protein) and the gene of interest, both under the control of the inducible *nitA* promoter. This design incorporates a positive feedback loop that increases inducibility (Fig S1a). On the other hand, the XylS/*Pm* system originated from the *Pseudomonas putida* TOL plasmid pWW0 and controls the expression of an operon encoding enzymes involved in aromatic hydrocarbon degradation (Gawin et al., 2017). Due to its applicability to a wide range of bacterial hosts, XylS/*Pm* is available in the SEVA repository (Martínez-García et al., 2023). The inducers needed for these promoters, ε-caprolactam and 3-methylbenzoate (P_nit_ and XylS/*Pm* system) showed to be non-toxic at the concentrations commonly used for gene expression (30 mM and 2 mM respectively, Fig S1b). These promoters were also cloned into the self-replicating vector pSEVA251 and transferred into *Synechocystis* by conjugation, including an empty vector as control. Transformed *Syn6803* were used to inoculate fresh cultures, which were grown for 24 hours before induction of expression by addition of ε-caprolactam or 3-methylbenzoate. The cells were cultivated for an additional 72 hours, and the fluorescence from expressed reporter (msfGFP) was determined (Fig. S1c). Any of those promoters display an increase of expression when the inducer was added to the culture. The signal for XylS/*Pm* was low; however, in P_nit_, it was ∼85 times higher than the control (cells harbouring the empty pSEVA yielded 0.6 rfu versus the 53.7 rfu for the cells harbouring the P_nit_ construction) suggesting a constitutive expression in *Syn6803*. Recently, it has been observed that promoter strength differs between strains probably due to several factors such as their GC content, their particularities in their gene expression, or their phylogenetic diversity (Trieu et al., 2026). Therefore, characterize unexplored inducible promoters from other microorganisms could be an important base for developing new genetic resources in cyanobacteria.

### Assembly of CRISPR-dCas9 into a pSEVA vector

CRISPRi technology reports applied to cyanobacteria can be found elsewhere (Huang et al., 2016; Higo et al., 2018; Knoot et al., 2020; Victoria et al., 2024). But so far, there has not been applied this edition tool using the SEVA modular technology. Among the advantages of using SEVA vectors are that i) they provide a reliable backbone for genetic operations across diverse hosts; ii) they count with a standardised Plug-and-Play architecture consisting of three interchangeable functional modules that can be swapped providing many different combinations without redesigning entire constructs; and that iii) they count with a broad assembly compatibility with traditional restriction-ligation as well as modern multi-part DNA assembly methods such as MoClo, Golden Gate, BioBricks, or BASIC assembly workflows (Haines et al., 2022; Martínez-García et al., 2023; de Lorenzo and Martinez-Garcia, 2025). Therefore, to improve CRISPR tools in cyanobacteria, dCas9 from *Streptococcus pyogenes* (Yao et al., 2016) was transferred into a pSEVA for CRISPRi-mediated gene repression. For *dCas9* expression, cLac promoters tested previously in this work were discarded for several reasons: i) cLac143: its leakiness could produce undesired repression in absence of IPTG and ii) cLac94: an 80-fold-change when IPTG is used could lead to toxic effects in the cell.

Riboswitches consist of two partially overlapping domains: an aptamer domain, which specifically binds an effector molecule, triggering a conformational change in the RNA structure, and an expression platform, which regulates gene expression by modulating transcription termination or translation initiation (Satta et al., 2023). The theophylline responsive riboswitch integrated into a rhamnose-inducible promoter system represents a dual control that provides a tight and tunable expression system in *Syn*6803 or *E. col*i (Liu et al., 2020; Della Valle et al., 2025). A tightly controlled CRISPR/Cas9 editing system for Syn6803, where the expression of Cas9, is controlled by a theophylline-inducible riboswitch has been built on the replicative pPMQAK1-T based-vector harboring RSF1010-replicon (Cengic et al., 2022).

For this work, we have chosen the system previously described to successfully control the expression of the DNase-dead Cpf1 nuclease in *Syn6803* in which a theophylline responsive riboswitch was integrated into the rhamnose-inducible promoter system to yield P_BADRSW_ promoter (Fig S2a) (Liu et al., 2020). This P_BADRSW_ promoter, which includes P_BAD_ promoter with a riboswitch, was selected to drive dCas9 expression in a newly built SEVA-CRISPRi vector. Previously, this promoter was tested in a pSEVA251; it showed low leakiness without inducers, and a 10-fold increase of fluorescence when rhamnose and theophylline were added to the culture (compared to uninduced cultures) (Fig S2b). Considering these results, a new SEVA was then constructed including dCas9 under P_BADRSW_ promoter in pSEVA351 (a plasmid with the same ori as pSEVA251).

On the other hand, to evaluate the CRISPRi-mediated repression, the eYFP was used as reporter, which was integrated into the *slr1181* neutral site of *Syn6803* genome under the constitutive P_c223_ promoter (Fig 2C). A guide sequence was designed for YFP knockdown targeting 32 nt from +1 transcription start. The sgRNA consisted of the guide sequence followed by dCas9 handle sequence and *S. pyogenes* terminator (Qi et al., 2013). The sgRNA_YFP_ was cloned in the same vector as dCas9 in pSEVA351 (Fig 2A, 2B).

**Fig. 2.**
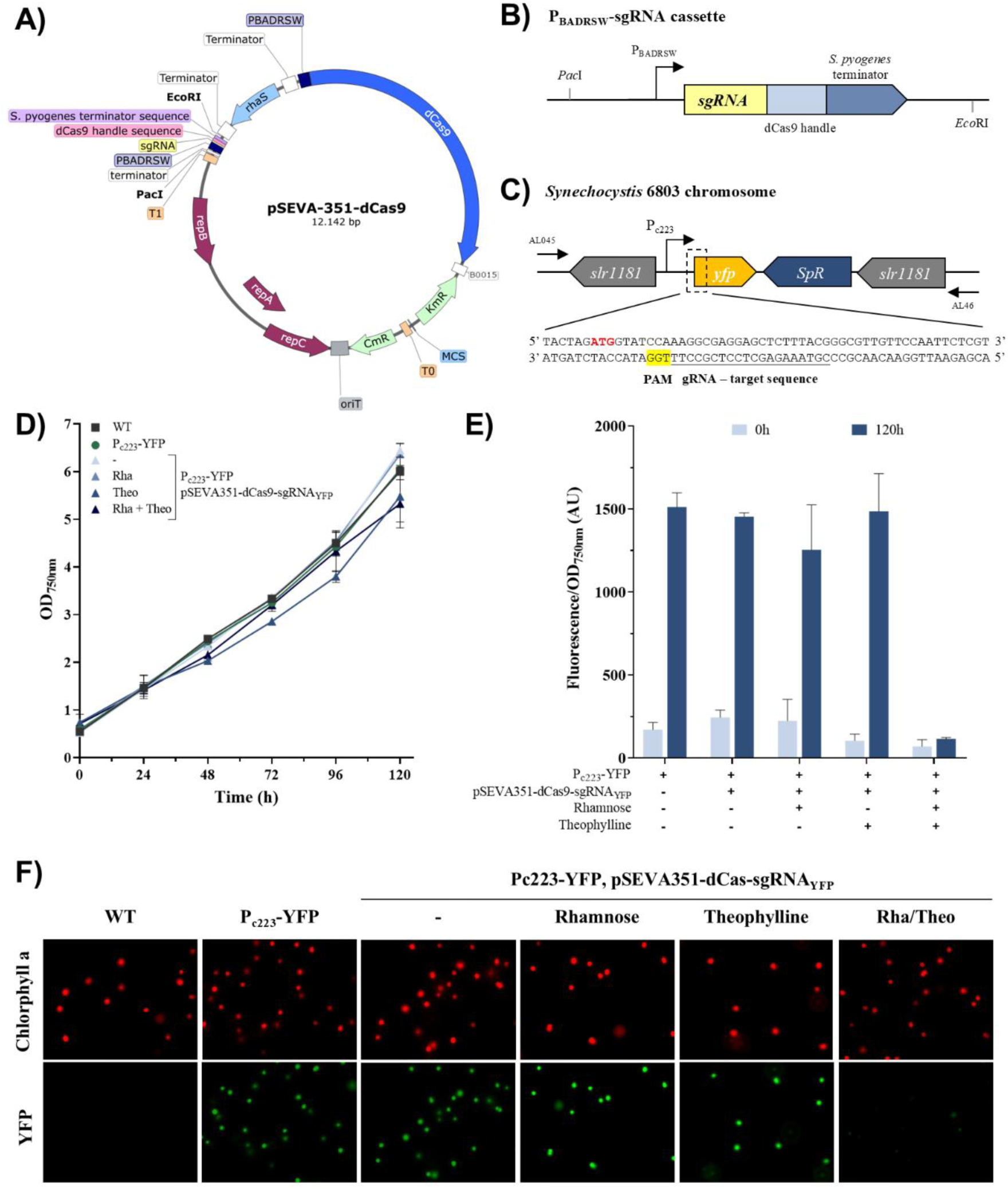
CRISPRi-mediated YFP repression in *Syn6803*with pSEVA351-dCas9-sgRNA_YFP_. **A)** Map of pSEVA351 with dCas9 expressed from P_BADRSW_ inducible promoter. **B)** Scheme of the sgRNA structure used for YFP repression. **C)** YFP integration into *Syn6803* chromosome in *slr1181* neutral site. **D)** Growth curves of *Syn6803* recombinants cultured in BG11 medium. Error bars represent the standard deviations observed from at least two independent experiments. **E)** Repression of reporter gene coding for eYFP by dCas9. pSEVA351-dCas9-sgRNA_YFP_ - refers to the uninduced strain. Rhamnose (Rha) and theophylline (Theo) were added to the culture with the concentrations 2 g/L and 2 mM respectively. Error bars represent the standard deviations observed from three independent experiments. **E)** Fluorescence microscopy of the CRISPRi-mediated repression assay 72 hours after induction. Chlorophyll a autofluorescence is shown as control.

Altogether, the repression of YFP, and then, a reduction in fluorescence, is expected in this construction when dCas9 is induced by rhamnose and theophylline.

### eYFP repression in *Syn6803* using pSEVA-dCas9

To test the functionality of the CRISPRi system in the eYFP-reporter *Syn6803*cells harboring the pSEVA-dCas9 vector, an induction assay was performed with rhamnose and theophylline. Optical density and fluorescence measurements were taken every 24 hours during a 4-day period. All strains showed a similar growth indicating that nor the presence of the pSEVA-dCas9 vector neither the inductors interfere with growth (Fig. 2d). Then we evaluated the fluorescence and initial values of fluorescence at time zero were quite similar among the different strains except in the pSEVA-dCas9 harboring strain under the presence of both inducers that yielded a 60% drop of fluorescence respect to the eYFP-reporter *Syn6803* control strain (Fig 2e).

Fluorescence after 72 hours growth was 8.8 higher respect to time zero in the control strain (Fig. 2e). The pSEVA-dCas9 harboring strain without the presence of both inducers or when it was grown just under the presence of rhamnose or theophylline yielded similar increases, raising the fluorescence level around 5.9, 5.6 or 14.2-fold respectively (Fig. 2e). The ratio of fluorescence 72h vs zero time was only 1.6-fold when both inducers were present.

When comparing the fluorescence values of the cas9 harboring strains compared to the control strain, a repression was only observed under the presence of rhamnose (a drop of 16.9%) or more drastically (a drop of 92.3%) under the presence of both theophylline and rhamnose after 72 hours growth.

This effect is also observed by fluorescence microscopy (Fig. 2f). Therefore, it is necessary to use both compounds (rhamnose and theophylline) to achieve complete repression. This can be explained because the riboswitch increases the strength and control of the promoter. It is also remarkable that theophylline alone has no effect on YFP repression (just a drop of fluorescence of 1.7% compared to the 16.9% drop in the presence of just rhamnose).

In other CrispRi experiments done in the model cyanobacterium *Syn*6803 in which Cas9 was used to repress green fluorescent protein (GFP), negligible levels were reported to be observed but when using putative aldehyde reductases or dehydrogenases, they just got a 50-95% repression (Yao et al., 2016). Therefore, further experiments with other reporters or genes should be done to check the pSEVA-dCas9 vector.

To check whether the repression observed was specific, as control, a guide sequence was designed for GFP knockdown (Fig. S3a). The sgRNA was selected to target the non-template strand and 28 nt downstream the start codon. The potential off-target binding sites for *yfp* had at least 8 mismatches, with 4 of them in the seed region. The designed sgRNA was cloned into pSEVA351-dCas9 and transformed into *Syn6803*expressing the YFP. When both inducers for the dCas9 harboring strains were used, the YFP fluorescence remained in the cell (97% of the fluorescence in the control), indicating that repression is occurring in a specific manner (Fig. S3b).

### Reversibility of YFP repression by SEVA-dCas9

Once that pSEVA-CRISPRi system was probed to be effective in inhibiting gene expression, the next step was to assess whether this repression was reversed by removing the inducers. For this, the previously induced culture was washed 3 times and resuspended in fresh BG11 without inducers at an initial OD_750_ of 0.8. Samples were taken at different times for fluorescence analysis (Fig. 3).

**Fig. 3.**
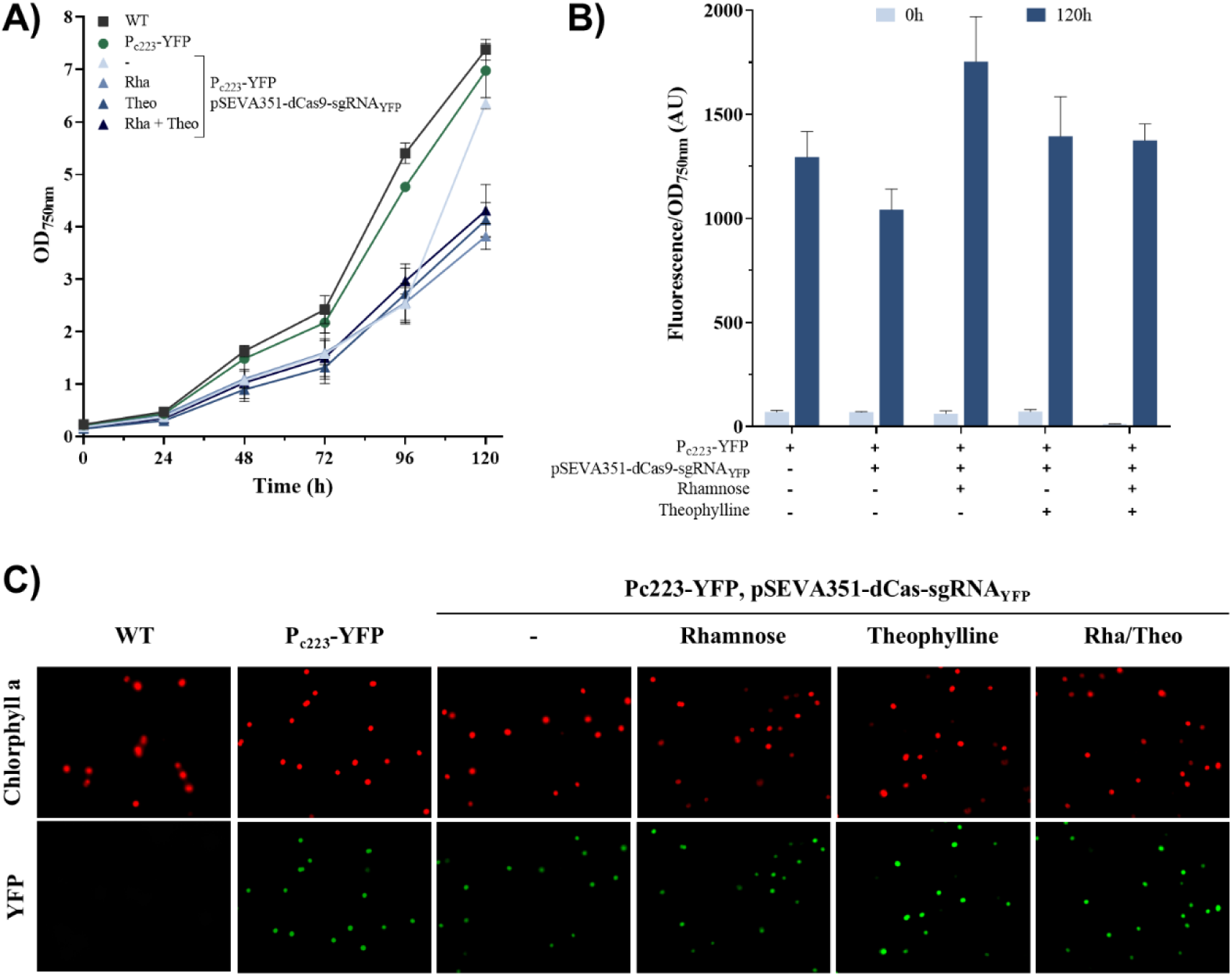
Recovery of the expression of YFP after release of the repression by the pSEVA351-dCas9. A) Growth curves of *Syn6803* strains cultured in fresh BG11 medium without inducers at an initial OD_750_ 0.8. Error bars represent the standard deviations observed from at least two independent experiments. B) Recovery of the reporter gene coding for YFP by removing inductors from the media. Fluorescence data was normalized to optical density at 750 nm. Error bars represent the standard deviations observed from three independent experiments. C) Fluorescence microscopy of the YFP expression 7 days after removing the inducers. Chlorophyll an autofluorescence is shown as control.

Initially, fluorescence values were quite similar in all samples except in the strain grown under the presence of both inducers that displayed a reduction of 84% respect to the YFP harboring strain control.

It was observed after 120h growth after retiring the presence of inducers, that fluorescence values were completely recovered in all strains, indicating that the repression was reversible.

## Conclusions

*Synechocystis* PCC6803 is an important model organism in scientific research and for industrial applications. Improving the genetic toolbox for *Syn6803* is a needed step for getting the most of its biotechnological potential. In this study different promoters on this strain have been evaluated. A low fluorescence signal was obtained for the constitutive promoter Pc223. Pnit and XylS/Pm promoters were leaky and yielded a high fluorescence signal in the absence of inducer (ε-caprolactam and 3-methylbenzoate respectively). The IPTG induced PcLac94 and PcLac143 promoters cloned into the self-replicating vector pSEVA251 using eYFP as reporter were the most effective ones: PcLac94 showed a low leakiness but higher induction rate (∼80-fold increase) while cLac143 was induced ∼25 times and yielded the highest fluorescence values after the induction. It is still needed to count on more effective inducible promoters.

On the other hand, in this study, a riboswitch-based, theophylline inducible CRISPRi/Cas9-system for easy use has been designed in Syn6803 based on seva351-plasmid. As a proof of concept, the suppression of eYFP expressed constitutively in Syn6803 was obtained only under the presence of the promoter activators in a reversible process. CRISPRi technology is still not widespread in cyanobacteria although it allows gene expression modification without editing the genome. The system presented here will help with the implementation of this technique on this strain.

## Availability of data and materials

All data generated or analysed during this study are included in this published article and its supplementary information files.

## Abbreviations

Cas: CRISPR associated protein
*Cm*^R^: chloramphenicol resistance gene
CRISPR: clustered regularly interspaced short palindromic repeats
eYFP: enhanced yellow fluorescent protein
GFP: Green Fluorescence Protein
IPTG: Isopropil ß-D-1-thiogalactopyranoside
*Km*^R^: kanamycin resistance gene
PAM: protospacer adjacent motif
PCR: polymerase chain reaction
SEVA: Standard European Vector Architecture
sgRNA: synthetic guide RNA
YFP: Yellow fluorescence protein.

## Acknowledgments

We are grateful to the Iberoamerican network Renuwal-Cyted 320RT0005 and Fotored UIU 2026 network.

## Author information

Govinda Guevara and Juana María Navarro have contributed equally to this work.

### Authors and Affiliations

Department of Biochemistry and Molecular Biology, c/Jose Antonio Novais 12, Universidad Complutense de Madrid, 28040 Madrid, Spain.

### Contributions

Co00nceptualization: GG, JMN, and SB. Writing and editing: SB, GG and JMN. Laboratory work and data analysis: SB, GG, AM and JMN. Project administration and funding acquisition: JMN. All authors have read and approved the final manuscript.

## Ethics declarations

### Ethics approval and consent to participate

Not applicable

### Consent for publication

Not applicable

## Additional information

Publisher’s note

Springer Nature remains neutral with regard to jurisdictional claims in published maps and institutional affiliations.

## Electronic supplementary material

**Table S1.**
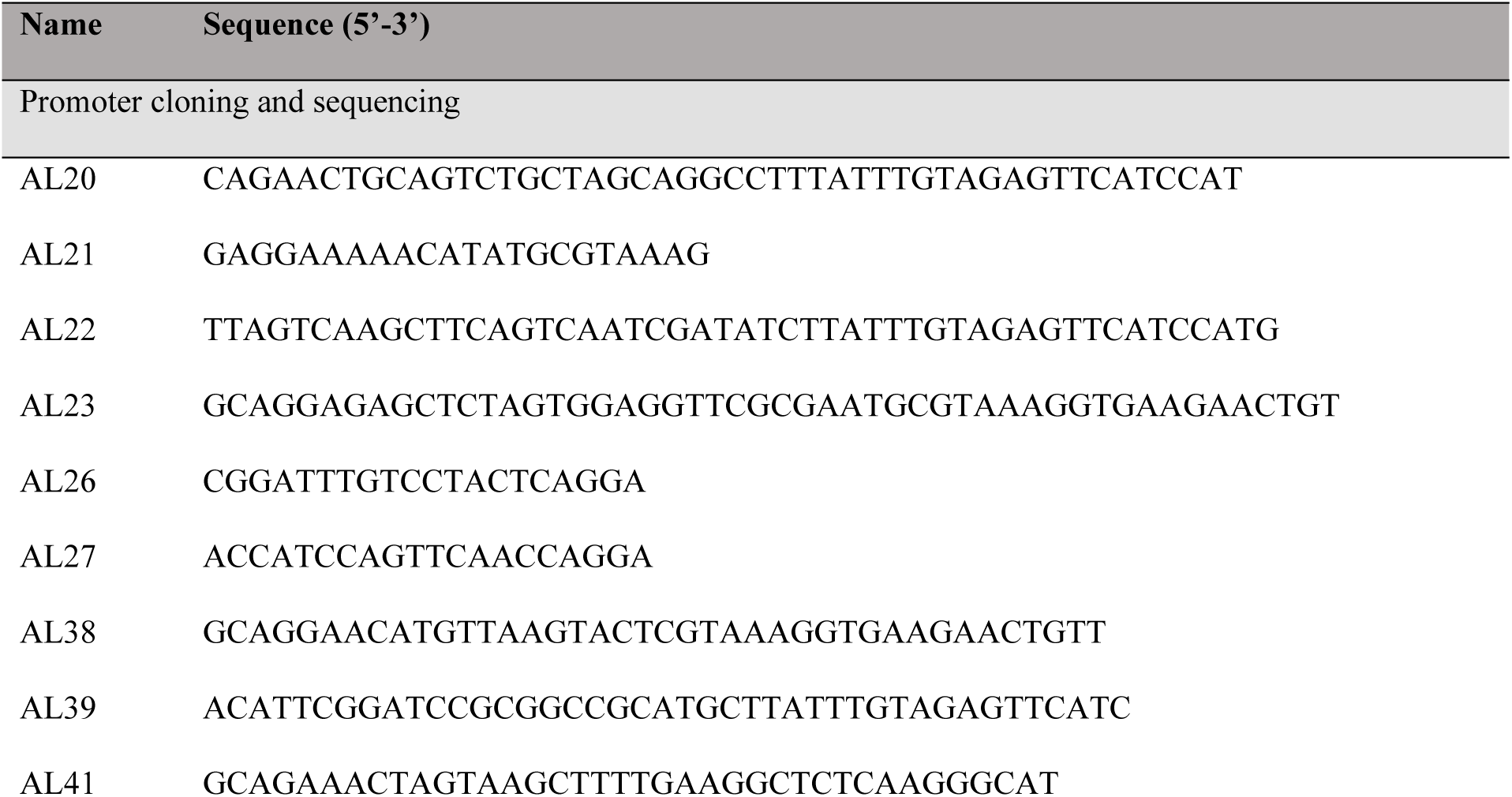
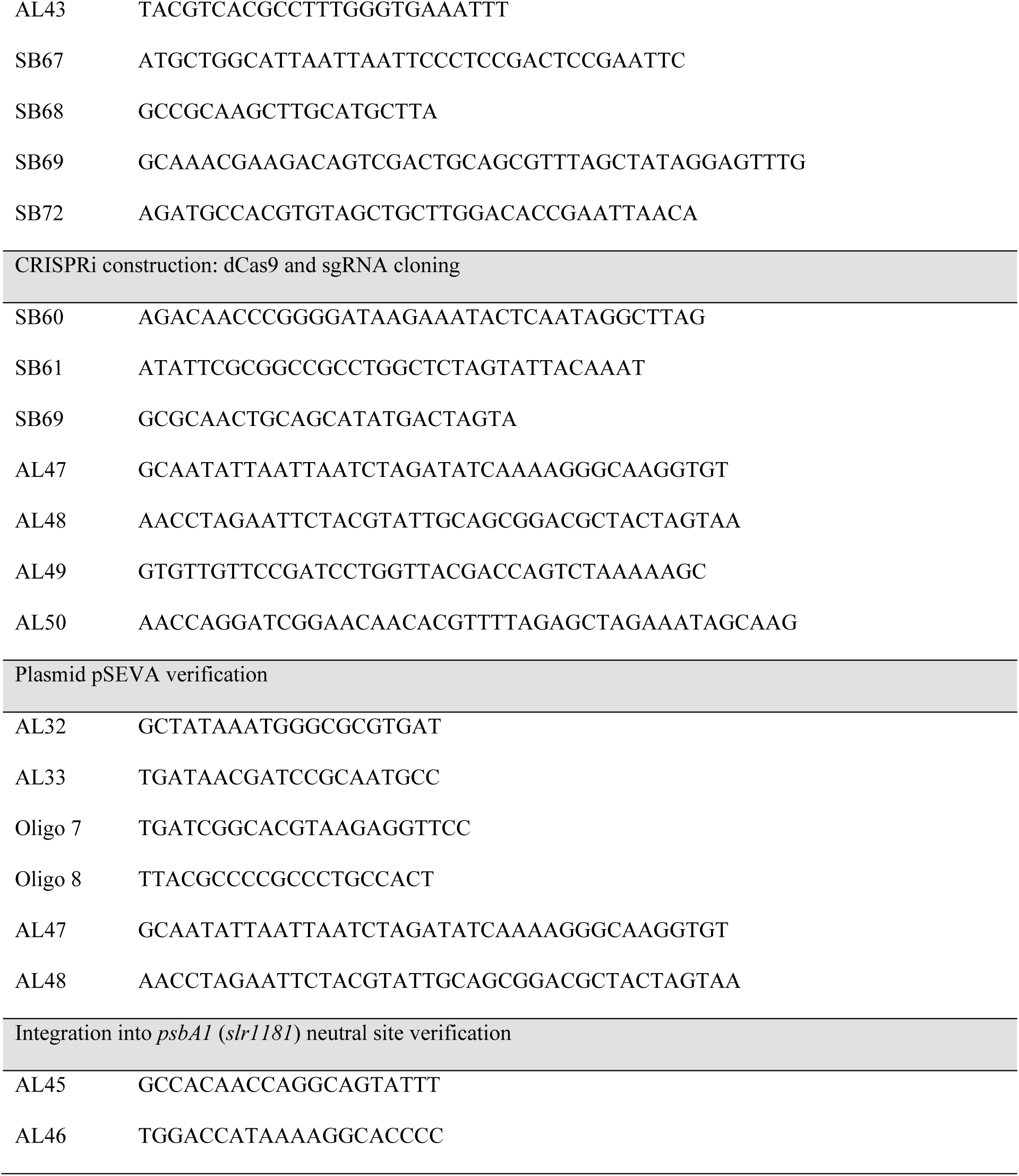
Primers used in this work.

**Table S2.**
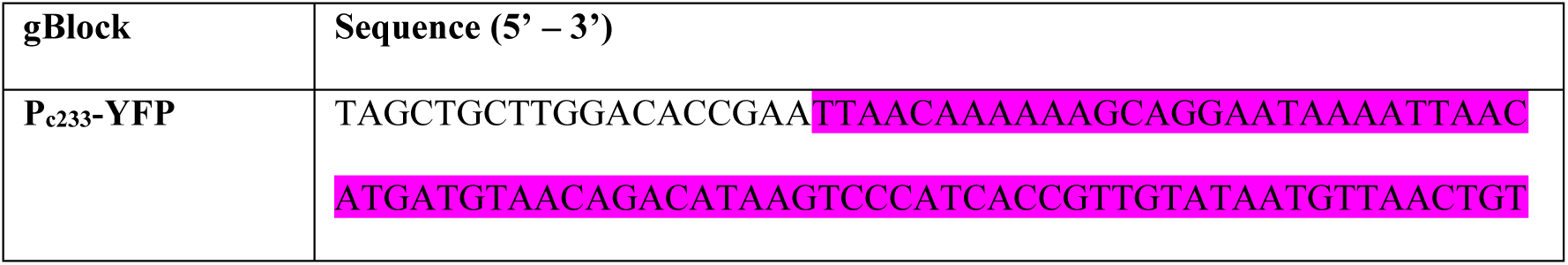
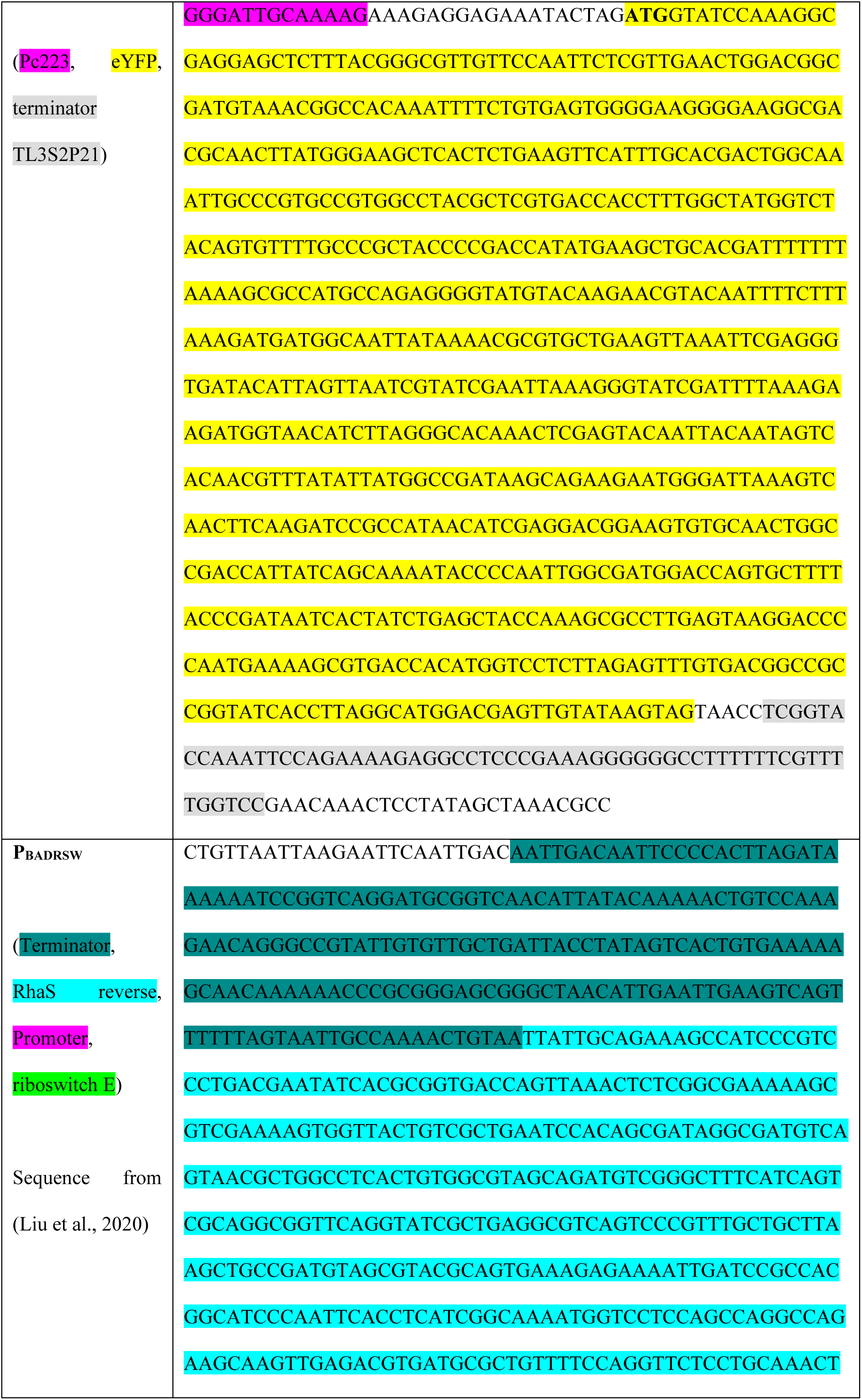
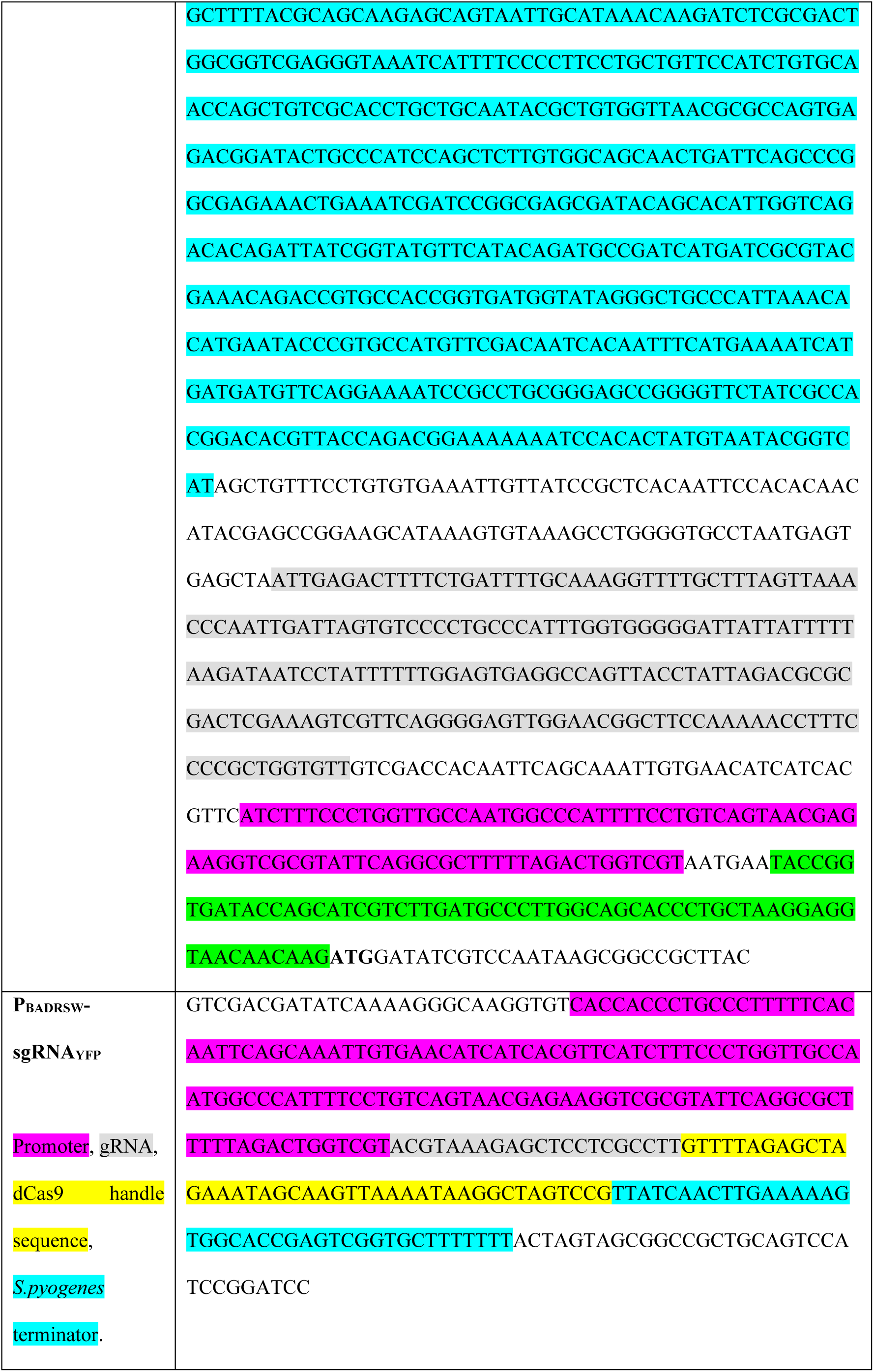
gBlocks used in this work.

TABLE S3. DATA AND STATISTICS

**Fig. S1.**
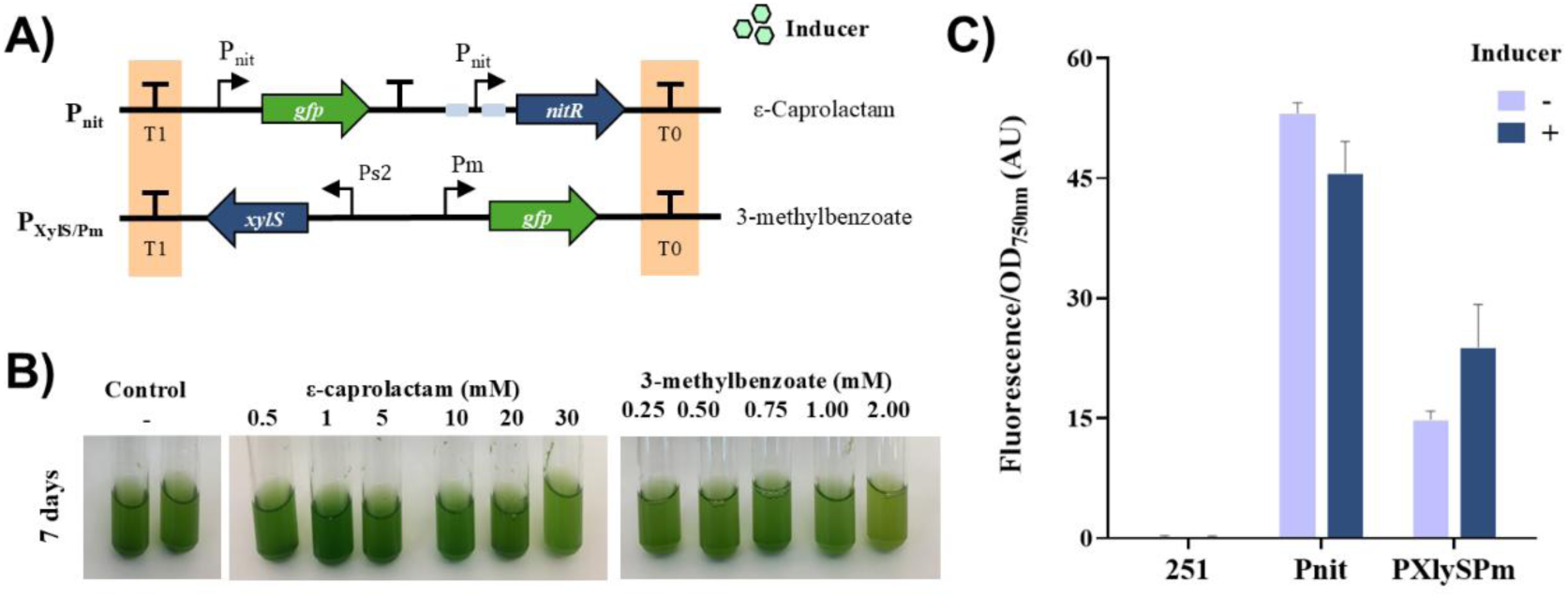
Inducible promoter activities in *Syn6803* from other bacteria. **A)** Diagram of promoters used on the replicating plasmid pSEVA251 in *Syn6803*. **B)** P_XylS/Pm_ induction. A *Synechocystis* culture was diluted to an OD_750nm_ of 0.8, and then the different conditions of inducers were added to BG11. As control, any compound was included in the culture. Pictures were taken 7 days of culture. **C)** Fluorescence measurements from the different promoters. Activities were measured as GFP fluorescence corrected by OD_750nm_ for cell cultures in BG11 medium. The results correspond to 72 hours post-induction, which was 20 mM of ε-caprolactam for P_nit_, and 1 mM of 3-methylbenzoate for P_XylS/Pm_. As control, a constitutive promoter was added. Error bars indicate standard deviations (*n*= 3).

**Fig. S2.**
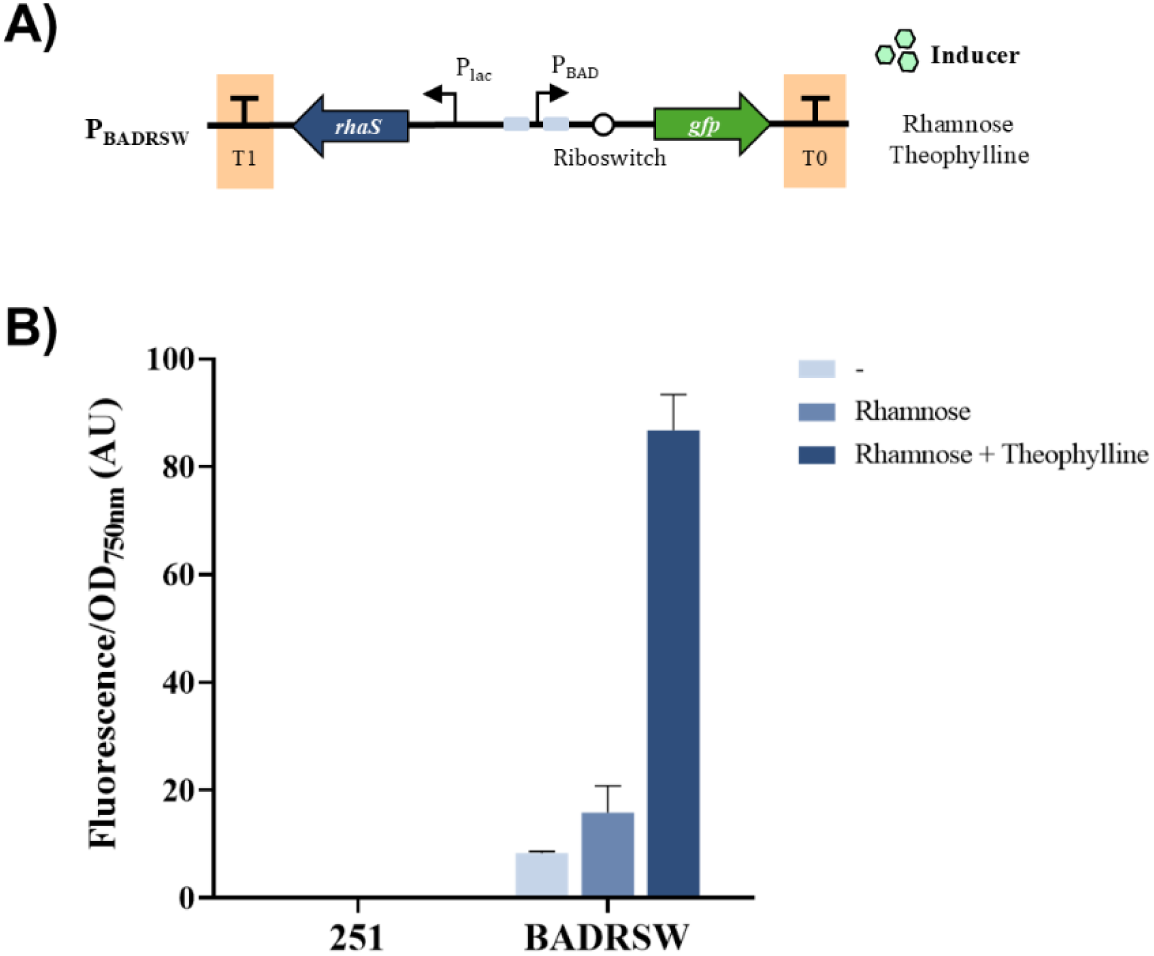
Validation of P_BADRSW_ promoter for msfGFP expression in pSEVA251. **A)** Diagram of P_BADRSW_ promoter on the replicating plasmid pSEVA251. **B)** Fluorescence measurements using different conditions of inducers. Results are shown as GFP fluorescence corrected by OD_750nm_ for cell cultures in BG11 medium. The results correspond to 72 hours post-induction with 2 g/L of rhamnose and/or 2 mM of theophylline. As control, pSEVA251 was added. Error bars indicate standard deviations (*n*= 3).

**Fig S3.**
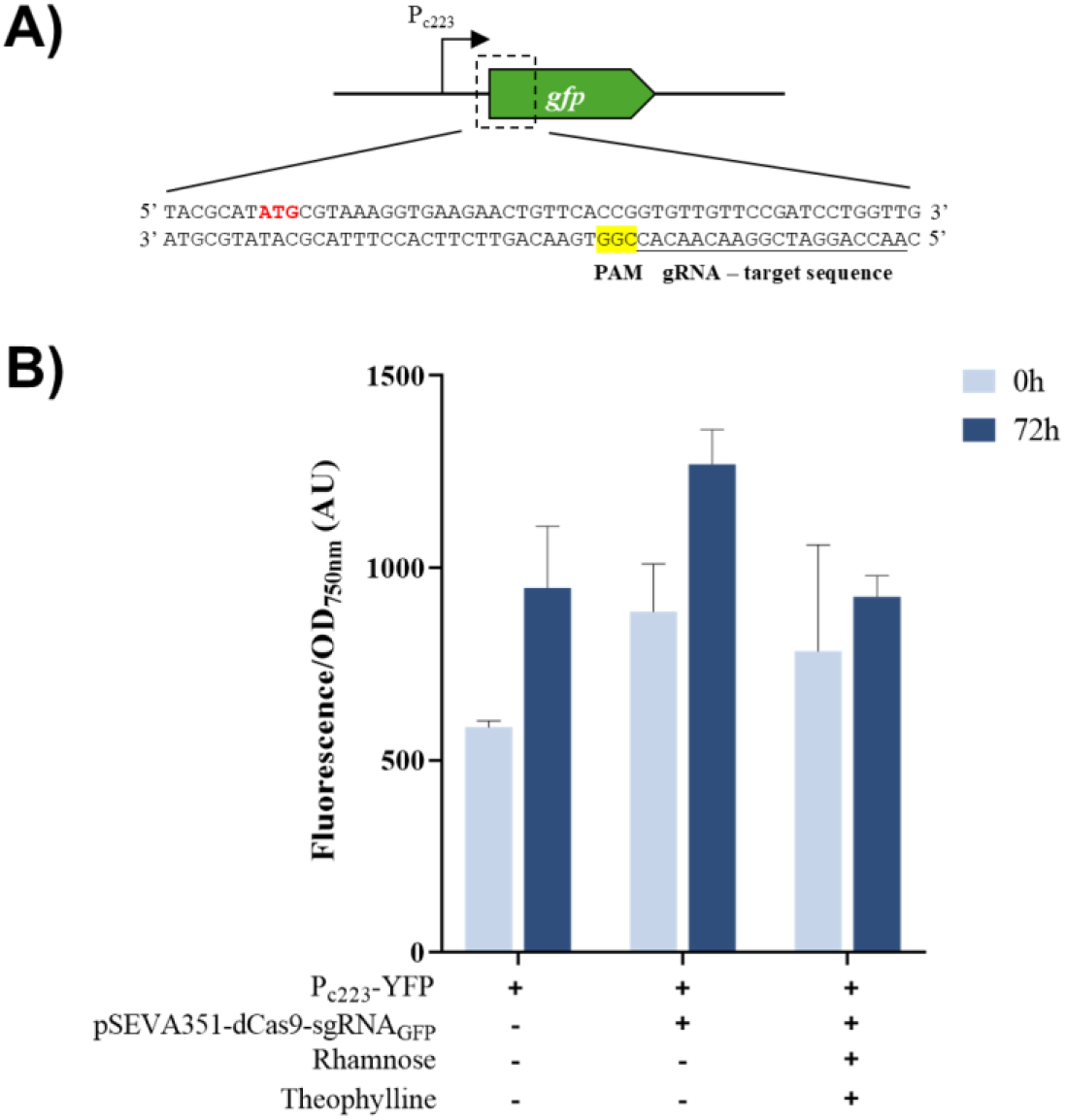
Specificity of pSEVA351-dCas9. **A)** Scheme of the sgRNA designed targeting GFP sequence. B) YFP fluorescence measurements using pSEVA351-dCAs9-sgRNA_GFP_. Rhamnose (Rha) and theophylline (Theo) were added to the culture with the concentrations 2 g/L and 2 mM respectively. Error bars represent the standard deviations observed from two independent experiments.

## REFERENCES

Albers, S.C. Peebles, C.A., 2017. Evaluating light-induced promoters for the control of heterologous gene expression in *Synechocystis* sp. PCC 6803. Biotechnology progress 33 (1), 45–53. 10.1002/btpr.2396.

Albers, S.C., Gallegos, V.A., Peebles, C.A., 2015. Engineering of genetic control tools in *Synechocystis* sp. PCC 6803 using rational design techniques. Journal of biotechnology 216, 36–46. 10.1016/j.jbiotec.2015.09.042.

Angeles, R., Carvalho, J., Hernandez-Martinez, I., Morales-Ibarria, M., Fradinho, J.C., Reis, M.A.M., Lebrero, R., 2025. Harnessing nature’s palette: Exploring photosynthetic pigments for sustainable biotechnology. N Biotechnol 85, 84–102. 10.1016/j.nbt.2025.01.001.

Baldanta, S., Guevara, G., Navarro-Llorens, J.M., 2022. SEVA-Cpf1, a CRISPR-Cas12a vector for genome editing in cyanobacteria. Microbial cell factories 21 (1), 103. 10.1186/s12934-022-01830-4.

Behle, A., Saake, P., Germann, A.T., Dienst, D., Axmann, I.M., 2020. Comparative dose-response analysis of inducible promoters in cyanobacteria. ACS synthetic biology 9 (4), 843–855. 10.1021/acssynbio.9b00505.

Bolay, P., Toepel, J., Buhler, B., 2025. Biotechnological applications of cyanobacteria: *Synechocystis* and *Synechococcus* strains. Advances in biochemical engineering/biotechnology 192, 155–191. 10.1007/10_2025_282.

Bourgade, B. Stensjö, K., 2022. Synthetic biology in marine cyanobacteria: Advances and challenges. Frontiers in microbiology 13, 994365. 10.3389/fmicb.2022.994365.

Cengic, I., Cañadas, I.C., Minton, N.P., Hudson, E.P., 2022. Inducible CRISPR/Cas9 allows for multiplexed and rapidly segregated single-target genome editing in *Synechocystis* Sp. PCC 6803. ACS synthetic biology 11 (9), 3100–3113. 10.1021/acssynbio.2c00375.

Chen, M., Dhakal, D., Eckhardt, C.W., Luesch, H., Ding, Y., 2025. Synthetic biology strategies for cyanobacterial systems to heterologously produce cyanobacterial natural products. Nat Prod Rep 42 (8), 1240–1250. 10.1039/d5np00009b.

Cheng, J., Zhang, K., Hou, Y., 2023. The current situations and limitations of genetic engineering in cyanobacteria: a mini review. Molecular biology reports 50 (6), 5481–5487. 10.1007/s11033-023-08456-8.

Das, M. Maiti, S.K., 2024. Employment of light-inducible promoter in genetically engineered cyanobacteria for photosynthetic isobutanol production with simulated diurnal sunlight and CO(2). Journal of biotechnology 393, 31–40. 10.1016/j.jbiotec.2024.07.014.

de Lorenzo, V. Martinez-Garcia, E., 2025. On the Choice of the Right Plasmid Vector(s) in the Times of Synthetic Biology. Microbial biotechnology 18 (12), e70273. 10.1111/1751-7915.70273.

Della Valle, S., Orsi, E., Creutzburg, S.C.A., Jansen, L.F.M., Pentari, E.N., Beisel, C.L., Steel, H., Nikel, P.I., Staals, R.H.J., Claassens, N.J., van der Oost, J., Huang, W.E., Patinios, C., 2025. Streamlined and efficient genome editing in *Cupriavidus necator* H16 using an optimised SIBR-Cas system. Trends in biotechnology 43 (6), 1470–1491. 10.1016/j.tibtech.2025.02.006.

Elhai, J. Wolk, C.P., 1988. Conjugal transfer of DNA to cyanobacteria. Methods Enzymol 167, 747–754. 10.1016/0076-6879(88)67086-8.

Elhai, J., Vepritskiy, A., Muro-Pastor, A.M., Flores, E., Wolk, C.P., 1997. Reduction of conjugal transfer efficiency by three restriction activities of *Anabaena* sp. strain PCC 7120. Journal of bacteriology 179 (6), 1998–2005. 10.1128/jb.179.6.1998-2005.1997.

Englund, E., Liang, F., Lindberg, P., 2016. Evaluation of promoters and ribosome binding sites for biotechnological applications in the unicellular cyanobacterium *Synechocystis* sp. PCC 6803. Scientific reports 6, 36640. 10.1038/srep36640.

Enkerlin, A.M., Hoffmann, U.A., Rapp, J., Miao, R., Postma, J., Feldbrugge, M., Link, H., Hudson, E.P., Selim, K.A., 2026. Deciphering the metabolic details of L-lysine toxicity in cyanobacteria. Plant Physiol 200 (4), 10.1093/plphys/kiag106.

Ferreira, E.A., Pacheco, C.C., Pinto, F., Pereira, J., Lamosa, P., Oliveira, P., Kirov, B., Jaramillo, A., Tamagnini, P., 2018. Expanding the toolbox for *Synechocystis* sp. PCC 6803: validation of replicative vectors and characterization of a novel set of promoters. Synth Biol (Oxf) 3 (1), ysy014. 10.1093/synbio/ysy014.

Gallego, I., Medic, N., Pedersen, J.S., Ramasamy, P.K., Robbens, J., Vereecke, E., Romeis, J., 2025. The microalgal sector in Europe: Towards a sustainable bioeconomy. N Biotechnol 86, 1–13. 10.1016/j.nbt.2025.01.002.

Gawin, A., Valla, S., Brautaset, T., 2017. The XylS/Pm regulator/promoter system and its use in fundamental studies of bacterial gene expression, recombinant protein production and metabolic engineering. Microbial biotechnology 10 (4), 702–718. 10.1111/1751-7915.12701.

Guevara, G., Olortegui Flores, Y., Fernández de Las Heras, L., Perera, J., Navarro Llorens, J.M., 2019. Metabolic engineering of *Rhodococcus ruber* Chol-4: A cell factory for testosterone production. PloS one 14 (7), e0220492. 10.1371/journal.pone.0220492.

Haines, M.C., Carling, B., Marshall, J., Shenshin, V.A., Baldwin, G.S., Freemont, P., Storch, M., 2022. basicsynbio and the BASIC SEVA collection: software and vectors for an established DNA assembly method. Synth Biol (Oxf) 7 (1), ysac023. 10.1093/synbio/ysac023.

Higo, A., Isu, A., Fukaya, Y., Ehira, S., Hisabori, T., 2018. Application of CRISPR interference for metabolic engineering of the heterocyst-forming multicellular cybacterium *Anabaena* sp. PCC 7120. Plant Cell Physiol 59 (1), 119–127. 10.1093/pcp/pcx166.

Hishida, A., Shirai, R., Higo, A., Matsutani, M., Nimura-Matsune, K., Takahashi, T., Watanabe, S., Ehira, S., Hihara, Y., 2024. CRISPRi knockdown of the *cyabrB1* gene induces the divergently transcribed *icfG* and *sll1783* operons related to carbon metabolism in the cyanobacterium *Synechocystis* sp. PCC 6803. The Journal of general and applied microbiology 70 (1), 10.2323/jgam.2024.01.001.

Hsu, P.D., Scott, D.A., Weinstein, J.A., Ran, F.A., Konermann, S., Agarwala, V., Li, Y., Fine, E.J., Wu, X., Shalem, O., Cradick, T.J., Marraffini, L.A., Bao, G., Zhang, F., 2013. DNA targeting specificity of RNA-guided Cas9 nucleases. Nature biotechnology 31 (9), 827–832. 10.1038/nbt.2647.

Huang, C.H., Shen, C.R., Li, H., Sung, L.Y., Wu, M.Y., Hu, Y.C., 2016. CRISPR interference (CRISPRi) for gene regulation and succinate production in cyanobacterium *S. elongatus* PCC 7942. Microbial cell factories 15 (1), 196. 10.1186/s12934-016-0595-3.

Jones, C.M., Parrish, S., Nielsen, D.R., 2021. Exploiting polyploidy for markerless and plasmid-free genome engineering in cyanobacteria. ACS synthetic biology 10 (9), 2371–2382. 10.1021/acssynbio.1c00269.

Kamalesan, A., Kumar, K.K., Nathan, B., Perumal, R., Natesan, S., Sampathrajan, V., 2026. Terpene-based biofuels production: a revolutionizing role of cyanobacteria through metabolic engineering, opportunities and challenges. Arch Microbiol 208 (11), 10.1007/s00203-026-05100-4.

Kelly, C.L., Taylor, G.M., Hitchcock, A., Torres-Méndez, A., Heap, J.T., 2018. A rhamnose-inducible system for precise and temporal control of gene expression in cyanobacteria. ACS synthetic biology 7 (4), 1056–1066. 10.1021/acssynbio.7b00435.

Kirtania, P., Hódi, B., Mallick, I., Vass, I.Z., Fehér, T., Vass, I., Kós, P.B., 2019. A single plasmid based CRISPR interference in *Synechocystis* 6803 - A proof of concept. PloS one 14 (11), e0225375. 10.1371/journal.pone.0225375.

Knoot, C.J., Biswas, S., Pakrasi, H.B., 2020. Tunable Repression of Key Photosynthetic Processes Using Cas12a CRISPR Interference in the Fast-Growing Cyanobacterium *Synechococcus* sp. UTEX 2973. ACS synthetic biology 9 (1), 132–143. 10.1021/acssynbio.9b00417.

Li, S., Sun, T., Liu, D., Zhang, T., Chen, L., Zhang, W., 2026. Iterative genome engineering platform enables efficient sucrose biosynthesis from CO(2) in photosynthetic Synechococcus elongatus UTEX 2973. Plant Biotechnol J 24 (10), 5284–5299. 10.1111/pbi.70702.

Lindberg, P., Kenkel, A., Bühler, K., 2023. Introduction to cyanobacteria. Advances in biochemical engineering/biotechnology 183, 1–24. 10.1007/10_2023_217.

Liu, D., Johnson, V.M., Pakrasi, H.B., 2020. A Reversibly induced CRISPRi system targeting photosystem II in the cyanobacterium *Synechocystis* sp. PCC 6803. ACS synthetic biology 9 (6), 1441–1449. 10.1021/acssynbio.0c00106.

Liu, X., Tang, K., Hu, J., 2024. Application of cyanobacteria as chassis cells in synthetic biology. Microorganisms 12 (7), 10.3390/microorganisms12071375.

Mager, M., Pineda Hernandez, H., Brandenburg, F., López-Maury, L., McCormick, A.J., Nürnberg, D.J., Orthwein, T., Russo, D.A., Victoria, A.J., Wang, X., Zedler, J.A.Z., Branco Dos Santos, F., Schmelling, N.M., 2023. Interlaboratory reproducibility in growth and reporter expression in the cyanobacterium *Synechocystis* sp. PCC 6803. ACS synthetic biology 12 (6), 1823–1835. 10.1021/acssynbio.3c00150.

Markley, A.L., Begemann, M.B., Clarke, R.E., Gordon, G.C., Pfleger, B.F., 2015. Synthetic biology toolbox for controlling gene expression in the cyanobacterium *Synechococcus* sp. strain PCC 7002. ACS synthetic biology 4 (5), 595–603. 10.1021/sb500260k.

Martínez-García, E., Fraile, S., Algar, E., Aparicio, T., Velázquez, E., Calles, B., Tas, H., Blázquez, B., Martín, B., Prieto, C., Sánchez-Sampedro, L., Nørholm, M.H.H., Volke, D.C., Wirth, N.T., Dvořák, P., Alejaldre, L., Grozinger, L., Crowther, M., Goñi-Moreno, A., Nikel, P.I., Nogales, J., de Lorenzo, V., 2023. SEVA 4.0: an update of the Standard European Vector Architecture database for advanced analysis and programming of bacterial phenotypes. Nucleic Acids Res 51 (D1), D1558–d1567. 10.1093/nar/gkac1059.

Meyer, A.J., Segall-Shapiro, T.H., Glassey, E., Zhang, J., Voigt, C.A., 2019. *Escherichia coli* "Marionette" strains with 12 highly optimized small-molecule sensors. Nature chemical biology 15 (2), 196–204. 10.1038/s41589-018-0168-3.

Miao, R., Jahn, M., Shabestary, K., Peltier, G., Hudson, E.P., 2023. CRISPR interference screens reveal growth-robustness tradeoffs in *Synechocystis* sp. PCC 6803 across growth conditions. The Plant cell 35 (11), 3937–3956. 10.1093/plcell/koad208.

Mills, L.A., McCormick, A.J., Lea-Smith, D.J., 2020. Current knowledge and recent advances in understanding metabolism of the model cyanobacterium *Synechocystis* sp. PCC 6803. Bioscience reports 40 (4), 10.1042/bsr20193325.

Murakami, H., Momokawa, N., Motomura, K., Kuroda, A., Hirota, R., 2026. Adaptive laboratory evolution optimizes an engineered phosphite utilization pathway in *Synechococcus elongatus* PCC 7942. J Biosci Bioeng 141 (3), 152–157. 10.1016/j.jbiosc.2025.11.006.

Pandey, A.K., Raman, S., Proff, R., Joshi, S., Kang, C.M., Rubin, E.J., Husson, R.N., Sassetti, C.M., 2009. Nitrile-inducible gene expression in mycobacteria. Tuberculosis (Edinburgh, Scotland) 89 (1), 12–16. 10.1016/j.tube.2008.07.007.

Pérez, A.A., Gajewski, J.P., Ferlez, B.H., Ludwig, M., Baker, C.S., Golbeck, J.H., Bryant, D.A., 2017. Zn2+-inducible expression platform for *Synechococcus* sp. strain PCC 7002 based on the smtA Promoter/Operator and smtB repressor. Applied and environmental microbiology 83 (3), 10.1128/aem.02491-16.

Qi, L.S., Larson, M.H., Gilbert, L.A., Doudna, J.A., Weissman, J.S., Arkin, A.P., Lim, W.A., 2013. Repurposing CRISPR as an RNA-guided platform for sequence-specific control of gene expression. Cell 152 (5), 1173–1183. 10.1016/j.cell.2013.02.022.

Satta, A., Esquirol, L., Ebert, B.E., 2023. Current Metabolic Engineering Strategies for Photosynthetic Bioproduction in Cyanobacteria. Microorganisms 11 (2), 10.3390/microorganisms11020455.

Seo, E., Choi, Y.N., Shin, Y.R., Kim, D., Lee, J.W., 2023. Design of synthetic promoters for cyanobacteria with generative deep-learning model. Nucleic Acids Res 51 (13), 7071–7082. 10.1093/nar/gkad451.

Taton, A., Ma, A.T., Ota, M., Golden, S.S., Golden, J.W., 2017. NOT gate genetic circuits to control gene expression in cyanobacteria. ACS synthetic biology 6 (12), 2175–2182. 10.1021/acssynbio.7b00203.

Taylor, G.M., Hitchcock, A., Heap, J.T., 2021. Combinatorial assembly platform enabling engineering of genetically stable metabolic pathways in cyanobacteria. Nucleic Acids Res 49 (21), e123. 10.1093/nar/gkab791.

Trieu, K.P., Bishe, B., Taton, A., Tieu, B.P., Golden, J.W., 2026. A set of constitutive promoters with graded strengths for gene expression in diverse cyanobacterial strains. Applied and environmental microbiology 92 (7), e0059826. 10.1128/aem.00598-26.

Vasudevan, R., Gale, G.A.R., Schiavon, A.A., Puzorjov, A., Malin, J., Gillespie, M.D., Vavitsas, K., Zulkower, V., Wang, B., Howe, C.J., Lea-Smith, D.J., McCormick, A.J., 2019. CyanoGate: a modular cloning suite for engineering cyanobacteria based on the plant MoClo Syntax. Plant Physiol 180 (1), 39–55. 10.1104/pp.18.01401.

Veit, M.C., Stauder, R., Bai, Y., Gabhrani, R., Schmidt, M., Klahn, S., Lai, B., 2025. The necessity of multi-parameter normalization in cyanobacterial research: A case study of the PsbU in *Synechocystis* sp. PCC 6803 using CRISPRi. J Biol Chem 301 (11), 110763. 10.1016/j.jbc.2025.110763.

Veit, M.C., Stauder, R., Bai, Y., Gabhrani, R., Schmidt, M., Klahn, S., Lai, B., 2026. Correction: The necessity of multi-parameter normalization in cyanobacterial research: A case study of the PsbU in Synechocystis sp. PCC 6803 using CRISPRi. J Biol Chem 302 (6), 111453. 10.1016/j.jbc.2026.111453.

Venero, O.M., Ndeh, R., Pan, A., Patrikainen, P., Eriksson, M., Akhgari, A., Kakko, L., Nedorezova, D., Mohamed, O., Thomson, L., Bonde, A., Kelanne, N., Aro, E.M., Adamala, K.P., Kallio, P., 2026. CyanoConstruct: simple platform for cyanobacterial expression construct assembly and translational tuning. BMC Biotechnol 26 (1), 10.1186/s12896-025-01098-7.

Victoria, A.J., Selao, T.T., Moreno-Cabezuelo, J.A., Mills, L.A., Gale, G.A.R., Lea-Smith, D.J., McCormick, A.J., 2024. A toolbox to engineer the highly productive cyanobacterium *Synechococcus* sp. PCC 11901. Plant Physiol 196 (2), 1674–1690. 10.1093/plphys/kiae261.

Walshe, J. Saha, S.K., 2026. Cyanobacteria as a photosynthetic chassis for metabolic pathway engineering with heterologous gene expression. Curr Issues Mol Biol 48 (6), 10.3390/cimb48060638.

Wang, B., Eckert, C., Maness, P.C., Yu, J., 2018. A Genetic toolbox for modulating the expression of heterologous genes in the cyanobacterium *Synechocystis* sp. PCC 6803. ACS synthetic biology 7 (1), 276–286. 10.1021/acssynbio.7b00297.

Wang, S.Y., Li, X., Wang, S.G., Xia, P.F., 2023. Base editing for reprogramming cyanobacterium *Synechococcus elongatus*. Metabolic engineering 75, 91–99. 10.1016/j.ymben.2022.11.005.

Włodarczyk, A., Selão, T.T., Norling, B., Nixon, P.J., 2020. Newly discovered *Synechococcus* sp. PCC 11901 is a robust cyanobacterial strain for high biomass production. Communications biology 3 (1), 215. 10.1038/s42003-020-0910-8.

Yao, L., Cengic, I., Anfelt, J., Hudson, E.P., 2016. Multiple gene repression in cyanobacteria using CRISPRi. ACS synthetic biology 5 (3), 207–212. 10.1021/acssynbio.5b00264.

Yao, L., Shabestary, K., Björk, S.M., Asplund-Samuelsson, J., Joensson, H.N., Jahn, M., Hudson, E.P., 2020. Pooled CRISPRi screening of the cyanobacterium *Synechocystis* sp PCC 6803 for enhanced industrial phenotypes. Nature communications 11 (1), 1666. 10.1038/s41467-020-15491-7.

Zhang, M., Luo, Q., Sun, H., Fritze, J., Luan, G., Lu, X., 2022. Engineering a controllable targeted protein degradation system and a derived OR-GATE-Type inducible gene expression system in *Synechococcus elongatus* PCC 7942. ACS synthetic biology 11 (1), 125–134. 10.1021/acssynbio.1c00226.

Zhang, P., Wang, H., Xu, H., Wei, L., Liu, L., Hu, Z., Wang, X., 2023. Deep flanking sequence engineering for efficient promoter design using DeepSEED. Nature communications 14 (1), 6309. 10.1038/s41467-023-41899-y.

Zhang, Y., Yang, S., Zheng, X., Tan, X., 2025. Cyanobacterial type I CRISPR-Cas systems: distribution, mechanisms, and genome editing applications. Front Bioeng Biotechnol 13, 1552030. 10.3389/fbioe.2025.1552030.

